# *OsHMA3* overexpression works more efficiently in generating low-Cd rice grain than *OsNramp5* knockout mutation

**DOI:** 10.1101/2024.07.11.603016

**Authors:** Yuejing Gui, Joanne Teo, Dongsheng Tian, Raji Mohan, Zhongchao Yin

## Abstract

Cadmium (Cd) is highly toxic and a carcinogen to humans. Rice is prone to absorbing Cd and accumulating it in the grain, which raises health concerns for rice consumers. OsNramp5 is a major transporter for Cd and manganese (Mn) uptake in rice, whereas OsHMA3 is a tonoplast-localized transporter for Cd detoxification. In this study, we compared the efficiency of *OsNramp5* knockout mutation and *OsHAM3* overexpression in reducing Cd content in the rice grain. The grain Cd content of the *OsNramp5* knockout mutants was significantly lower than that of the wild-type rice T5105. However, the *OsNramp5* knockout mutants still had much higher grain Cd content than the similar *OsNramp5* mutants reported previously or the *OsHAM3* overexpression line developed in our previous study. Pyramiding the *OsNramp5* mutant allele and the *OsHAM3* transgene in a double homozygous line could not further reduce grain Cd content. The *OsNramp5* gene in T5105 has a haplotype II promoter, and its knockout mutation partially impairs Mn uptake in rice. Our results demonstrate that *OsHMA3* overexpression works more efficiently in generating low-Cd rice grain than *OsNramp5* knockout mutation without affecting Mn uptake in rice.

## INTRODUCTION

Cadmium (Cd) and its compounds are highly toxic for most living organisms. Long-term exposure of the human body to Cd may lead to cancer and organ system toxicity. Rice is a staple cereal crop for over 40% of the world’s population. Compared to other crops, rice is prone to absorbing Cd and accumulating it in grain, which makes rice the major contributor to dietary Cd exposure. Several genes on Cd uptake, transport, and detoxification in rice have been isolated and characterized in recent years (Ueno et al., 2010; Miyadate et al., 2011; Ishikawa et al., 2012; Ishimaru et al., 2012; Sasaki et al., 2012). Among these, OsNramp5 is a major Cd transporter in rice (Ishikawa et al., 2012; Ishimaru et al., 2012; Sasaki et al., 2012). The *OsNramp5* knockout mutant significantly reduces Cd content in the rice grain (Ishikawa et al., 2012). The *OsNramp5* knockout mutants derived from gene editing were used to generate low-Cd rice (Tang et al., 2017; Hu et al., 2024). Since OsNramp5 is also a major transporter for manganese (Mn) uptake in rice and Mn is essential for rice growth and development, the *OsNramp5* knockout mutation might result in impaired growth, development, reproduction and stress response, especially when grown at low Mn concentrations (Sasaki et al., 2012; Dong et al., 2021). OsHMA3 is a tonoplast-localized transporter that transports Cd from the cytosol into vacuoles for Cd detoxification, which effectively inhibits Cd loading into the xylem for long-distance transport to the shoot and the grain (Ueno et al., 2010; Miyadate et al., 2011). Overexpression of the *OsHMA3* gene in rice significantly reduced Cd accumulation in the grain and enhanced Cd tolerance (Ueno et al., 2010; Sasaki et al., 2014; Shao et al., 2018). Following a similar approach, we recently produced low-Cd rice lines in the genetic background of rice variety T5105 by overexpressing *OsHMA3* under the control of rice *OsActin1* promoter (Gui et al., 2024). In this study, we generated *OsNramp5* knockout mutants using CRISPR/Cas9 technology and compared them with the *OsHAM3* overexpression lines on their efficiency in reducing Cd content in rice grain.

## MATERIALS AND METHODS

### Plant materials, growth conditions, and Cd treatment

T5105 is an improved aromatic rice line in KDML105 genetic background (Luo and Yin, 2013). Nipponbare is a japonica cultivar. HMA3-L3 is an *OsNramp5* overexpression line carrying the *P*_*Actin1*_*:cHMA3:T*_*Nos*_ gene in T5105 genetic background developed in the previous study (Gui et al., 2024). NH is a double homozygous rice line in T5105 genetic background harbouring the *OsNramp5* mutant allele from *nramp5-L45* and the *P*_*Actin1*_*:cHMA3:T*_*Nos*_ gene from HMA3-L3. Rice plants were grown in pot soil in a greenhouse at 24 - 33 °C under natural light with a day length of 12 to 12.10 hours. The soil used for rice plantation was a 1:1 mixture (by weight) of topsoil and BVB Peatmoss (Kekkilä-BVB), containing background levels of Cd at 0.436 mg/kg and Mn at 734.774 mg/kg, respectively. For the Cd treatment, the control mixed soil was supplemented with 3 mg/kg of Cd supplied as CdSO_4_ (Gui et al., 2024).

### Constructs and rice transformation

Gene editing on *OsNramp5* was carried out according to the method described previously (Tang et al., 2017). A specific DNA target sequence and its adjacent protospacer adjacent motif (PAM) (AGGTTCTTCCTGTACGAGAGCGGG) were designed for CRISPR/Cas9-mediated gene editing of exon 9 in *OsNramp5*. The expression cassette, containing the target sequence and the gRNA under the rice snRNA U3 promoter in vector pYLsgRNA-OsU3, was amplified and cloned into the vector pYLCRISPR/Cas9P_Ubi_-H to make binary construct pYLCas9-Nramp5. pYLCas9-Nramp5 was introduced into the *Agrobacterium tumefaciens* strain AGL1 by electroporation. The *Agrobacterium*-mediated transformation of T5105 was performed according to the procedures described previously (Gui et al., 2024).

### ICP-MS

The contents of Cd and Mn in the de-husked but unpolished brown rice were determined by ICP-MS (7700S, Agilent Technologies, USA) as described in the previous study (Gui et al., 2024).

### qRT-PCR

Total RNA extraction from rice root tissues, the first-strand cDNA synthesis and qRT-PCR analysis were carried out according to the methods described in the previous study (Gui et al., 2024). The qRT-PCR results were normalized against the rice elongation factor gene *OsEF-1α* (Os03g0178000). The relative expression levels of *OsNramp5* were determined using the 2^−ΔΔ*C*t^ relative quantification method with the expression level in Nipponbare arbitrarily set to 1. The oligo DNA primer pairs are 5’GTCGCCGTCGTCTACCT3’/5’GTACGGCAAGGGCTCGT3’ for the *OsNramp5* gene and 5’GCACGCTCTTCTTGCTTTC3’/5’AGGGAATCTTGTCAGGGTTG3’ for the *OsEF-1α* gene.

## RESULTS AND DISCUSSION

We recently developed low-Cd rice lines in the genetic background of rice variety T5105 by overexpressing *OsHMA3* under the control of rice *OsActin1* promoter (Gui et al., 2024). To suppress Cd uptake, we also generated three independent knockout mutants of *OsNramp5* by using CRISPR-Cas9 technology (Fig. 1A and 1B). The Cd contents in the grain of the *OsNramp5* knockout mutants (0.678±0.336 mg/kg to 0.715±0.217 mg/kg) were significantly lower than that of T5105 (1.561±0.398 mg/kg) when they were grown in Cd-contaminated soil (Fig. 1C). However, the Cd content in the grain of the *OsNramp5* knockout mutants was still significantly higher than that in the grain of the *OsNramp5* knockout mutants obtained through mutagenesis or gene editing in the previous reports (Ishikawa et al., 2012; Tang et al., 2017). As for the efficiency in grain Cd reduction, the Cd content in the grain of the *OsNramp5* knockout mutants was partially reduced by 54.2% to 56.6% over that of T5105, which was also significantly lower than that in the previous reports (Ishikawa et al., 2012; Tang et al., 2017). Like the *OsNramp5* mutants reported in the previous studies (Sasaki et al., 2012), the knockout mutation of the *OsNramp5* gene in T5105 partially affected Mn uptake and resulted in lower grain Mn content than the wild-type rice when they were grown in either control or Cd-contaminated soil (Fig. 1D). However, no significant change in the phenotype or seed setting rate was observed with the *OsNramp5* knockout mutants, indicating that the partial reduction in Mn uptake did not affect plant growth and development, possibly due to the high background level of Mn concentration at 734.774 mg/kg in the control soil (Sasaki et al., 2012).

**Fig. 1.**
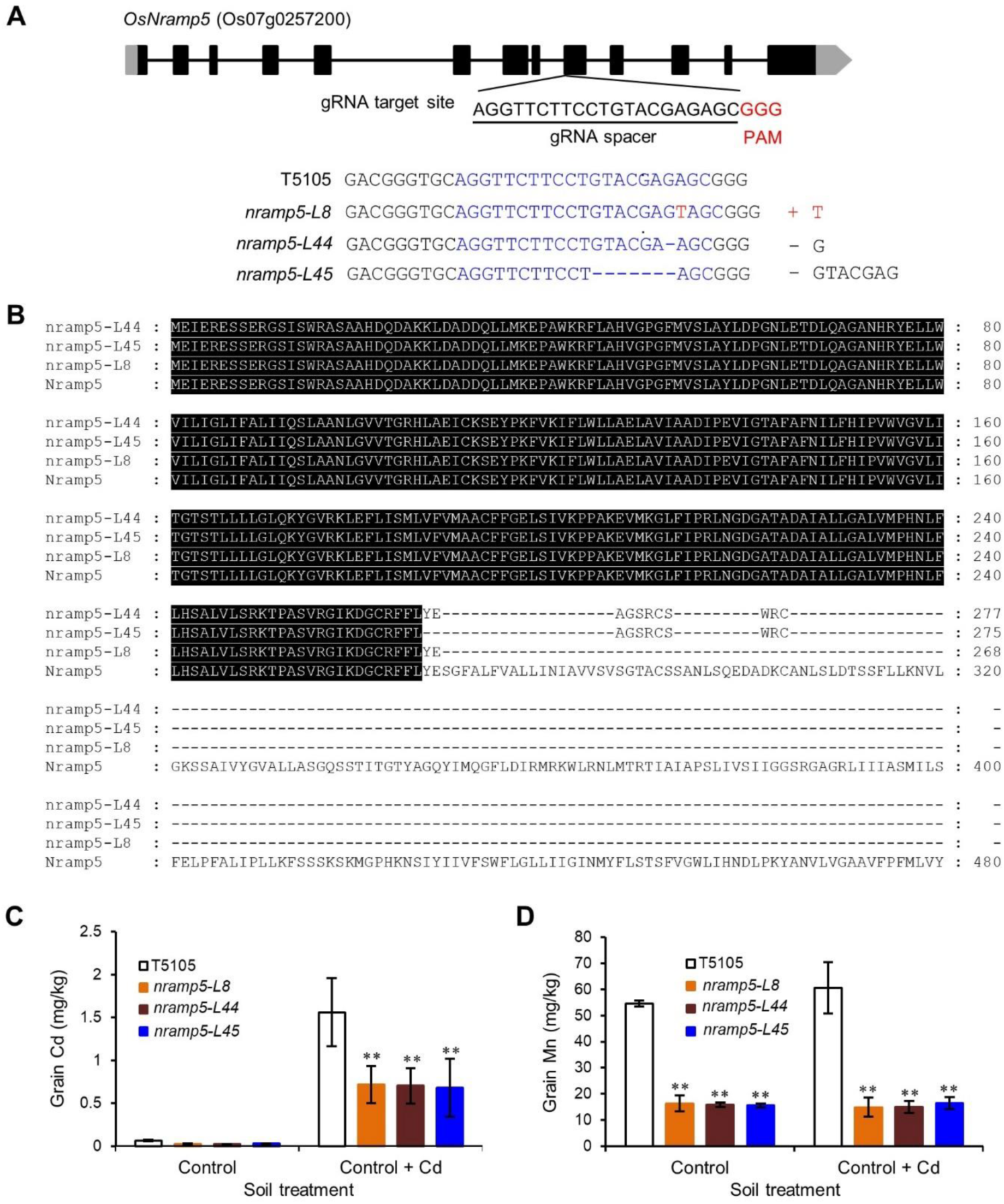
Generation of *OsNramp5* knockout mutants. (A) Schematic diagram of *OsNramp5* gene structure. UTRs, exons, and introns are indicated by grey blocks, black blocks, and lines, respectively. The nucleotide sequences of the CRISPR/Cas9 target together with PAM (GGG) and the gene edited *OsNramp5* mutants are shown at the bottom of the gene structure. Deletions and insertions are indicated by dashes and red letters, respectively. (B) Protein alignment of OsNramp5 and its mutants. The amino acid sequences were aligned by Clustral W method under the software MegAlign and viewed by software GeneDoc. The identical residues among all aligned proteins were highlighted in black. (C) Grain Cd content of T5105 and *OsNramp5* knockout mutants. (D) Grain Mn content of T5105 and *OsNramp5* knockout mutants. Asterisks in (C) and (D) indicate a significant difference between T5105 and *OsNramp5* mutants at ** *P* < 0.01 by Student’s *t*-test. All data are means ± SD of at least three biological replicates.

The representative *OsNramp5* mutant line *nramp5-L45* was then selected for a comparison study with HMA3-L3, a representative *OsHMA3* overexpression line in the previous study (Gui et al., 2024). The Cd content in the grain of *nramp5-L45* was 1.038±0.079 mg/kg when it was grown in Cd-contaminated soil, which was lower than that of T5105 at 1.720±0.137 mg/kg, but still much higher than the maximum level (ML) of Cd in rice at 0.4 mg/kg set by the Codex Alimentarius Commission (Codex) (Fig. 2A) (Commission, 2006). Pyramiding the *OsNramp5* mutant allele from *nramp5-L45* and the *P*_*Actin1*_*:cHMA3:T*_*Nos*_ gene from HMA3-L3 in a double homozygous line, designated as NH, did not result in any further significant Cd reduction in the grain of NH if compared to that of HMA3-L3 (Fig. 2A). The grain Cd contents of HMA3-L3 and NH were 0.027±0.005 mg/kg and 0.020±0.003 mg/kg, respectively (Fig. 2A). Both were significantly lower than the set ML of Cd in rice by Codex. Both *nramp5-L45* and NH had lower grain Mn contents than T5105 (Fig.2B). However, there was no significant difference in Mn content in the grain between HMA3-L3 and T5105, indicating that, unlike the *OsNramp5* knockout mutation, the *OsHMA3* overexpression did not affect Mn uptake in rice (Fig. 2B) (Gui et al., 2024). The partial impairment of Cd and Mn accumulation in the grain of the *OsNramp5* knockout mutants also suggests that, in addition to OsNramp5, the rice should have other transporters for Cd and Mn uptake. Indeed, the rice transporter OsNramp1 was found to contribute to Cd and Mn uptake (Takahashi et al., 2011; Chang et al., 2020). The two rice metal transporters have redundancy in Cd and Mn uptake.

**Fig. 2.**
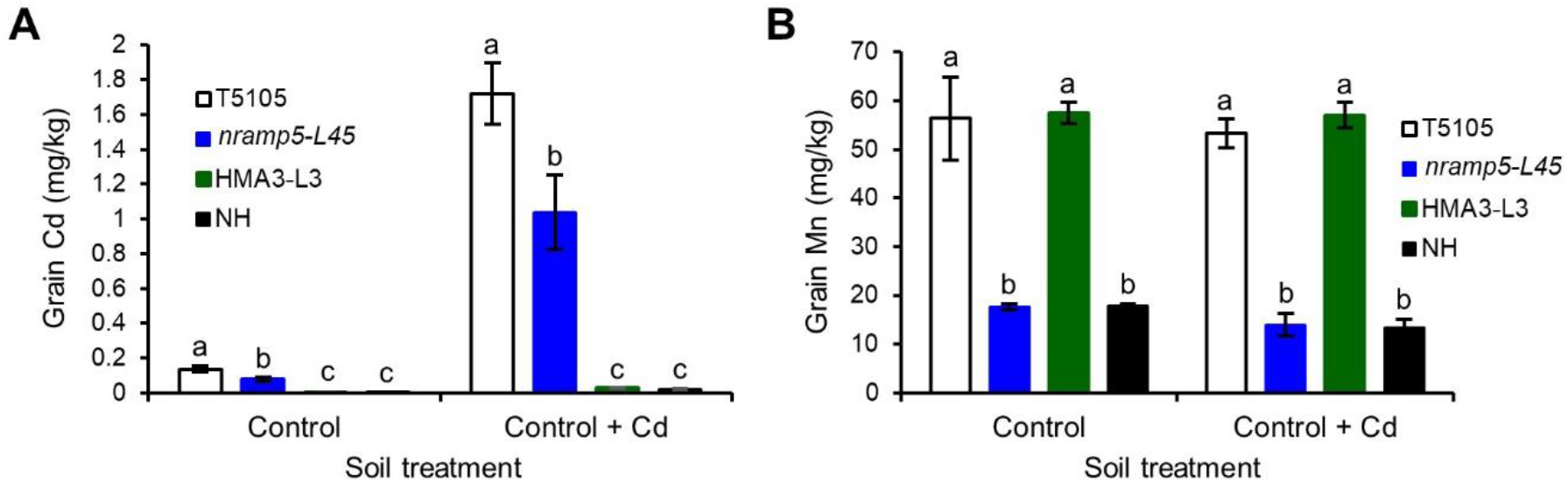
Cd and Mn contents in the grain of rice lines. (A) Cd content in the grain of T5105, *nramp5-L45*, HMA3-L3 and NH. (B) Mn content in the grain of T5105, *nramp5-L45*, HMA3-L3 and NH. Different letters in (A) and (B) indicate statistically different at *P* < 0.05 according to LSD’s test. All data are means ± SD of at least three biological replicates.

Based on the single nucleotide polymorphism (SNP) in the promoter, the *OsNramp5* genes in different rice varieties could be classified into three haplotypes, Haplotype I, Haplotype II and Haplotype III (Liu et al., 2017). *OsNramp5* carrying a Haplotype I promoter has a higher expression level than *OsNramp5* carrying a Haplotype II or Haplotype III promoter (Liu et al., 2017). Rice varieties carrying *OsNramp5* a Haplotype I promoter also accumulate more Mn in brown rice (Liu et al., 2017). Rice genome sequencing revealed that the *OsNramp5* gene in T5105 carries a Haplotype II promoter (Fig. 3A) (Gui et al., 2024). Further qRT-PCR analysis confirmed that the *OsNramp5* gene in T5105 had a lower expression level in root than the *OsNramp5* gene in Nipponbare, a cultivar harbouring an *OsNramp5* gene carrying a Haplotype I promoter (Fig. 3B) (Liu et al., 2017). On the one hand, due to the redundancy between *OsNramp1* and *OsNramp5*, the relatively low expression level of the *OsNramp5* gene in T5105 might weaken the contribution of *OsNramp5* to Cd and Mn uptake in rice. Therefore, the single knockout mutation of *OsNramp5* may only partially impairs Cd and Mn uptake in rice. On the other hand, as Mn is essential for rice, *OsNramp5* knockout mutants may affect rice growth, development, yield and tolerance to high temperature under the condition of insufficient Mn supply from soil (Sasaki et al., 2012; Dong et al., 2021).

**Fig. 3.**
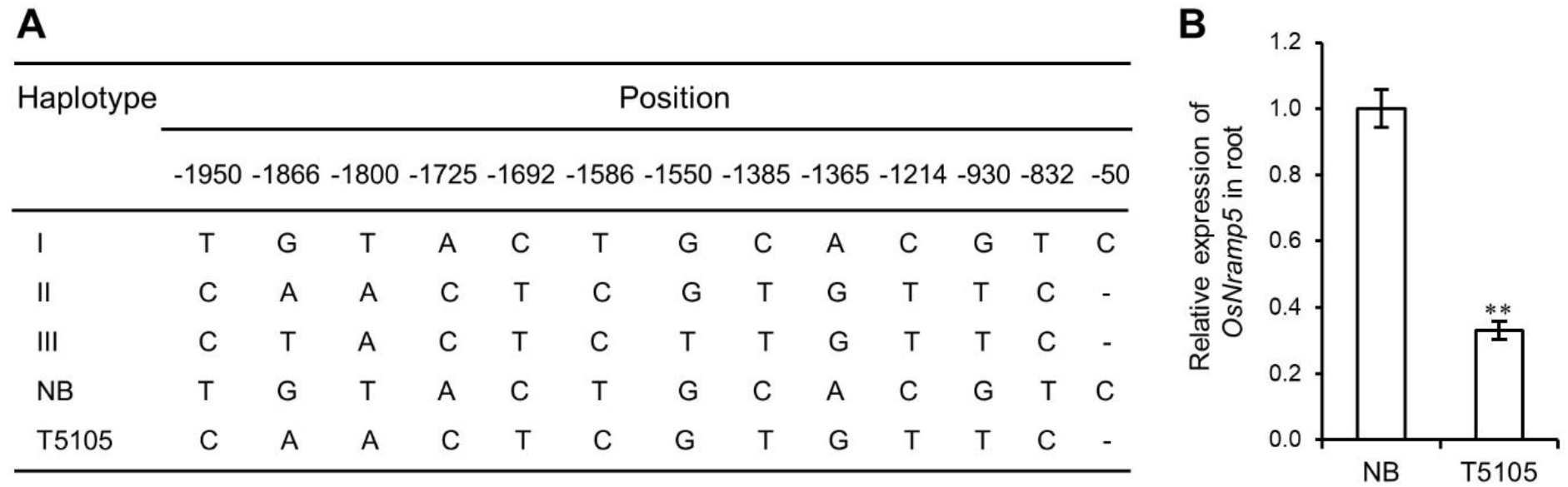
Haplotypes of *OsNramp5* promoters in rice and their relative expression in root. (A) Haplotypes of *OsNramp5* promoters and SNPs. The SNPs in Haplotype I to Haplotype III promoters of *OsNramp5* genes published by Liu et al., (2017) were listed as the references above the SNPs in the promoters of the *OsNramp5* genes in Nipponbare (NB) and T5105. (B) Relative expression of the *OsNramp5* genes in the root of Nipponbare and T5105 detected by qRT-PCR. The expression level of *OsNramp5* in the root of Nipponbare was set as 1. Asterisks in (B) indicate a significant difference between Nipponbare and T5105 at ** *P* < 0.01 by Student’s *t*-test.

Based on the results in this study and taking into consideration the above-mentioned concerns, we conclude that the *OsHMA3* overexpression approach works more efficiently in generating low-Cd rice grain than the *OsNramp5* knockout approach without affecting Mn uptake in rice.

## LIMITATIONS

The contribution of *OsNramp5* to Cd and Mn uptake and transport may work in a haplotype- or variety-dependent manner. The conclusion in this report was drawn based on results obtained from the study with variety T5105. It remains to be further investigated whether this also applies to other rice varieties or cultivars.

## DECLARATION

## Acknowledgements

The authors thank Prof. Yao-Guang Liu for providing gene editing constructs pYLsgRNA-OsU3 and pYL-CRISPR/Cas9P_Ubi_-H.

## Funding

This work was supported by intramural research funds for Temasek Life Sciences Laboratory from Temasek Trust and a fund from the National Research Foundation (NRF), Prime Minister’s Office, Singapore, on the Disruptive & Sustainable Technology for Agricultural Precision (DiSTAP).

## Author’s contributions

YZ conceived the research. GY, TJ and TD conducted the experiments. YZ, GY, TD and RM wrote the article.

## Availability of data and material

The *OsNramp5* knockout mutants developed in this study are available for non-commercial research purposes based on signing a Material Transfer Agreement defined by the Intellectual Property Office of Temasek Life Sciences Laboratory Ptd, Singapore.

## Competing interests

Temasek Life Sciences Laboratory Ltd has filed a patent on the method of generating low-As and low-Cd rice grains with ZY and YG as the inventors (PCT APPLICATION NO. PCT/SG2023/050847). The *OsHMA3* overexpression line HMA3-L3 used in this study was included in the patent.

## Notes

### Summary of Updates

Cadmium (Cd) is highly toxic and a carcinogen to humans. Rice is prone to absorbing Cd and accumulating it in the grain, which raises health concerns for rice consumers. OsNramp5 is a major transporter for Cd and manganese (Mn) uptake in rice, whereas OsHMA3 is a tonoplast-localized transporter for Cd detoxification. In this study, we compared the efficiency of OsNramp5 knockout mutation and OsHAM3 overexpression in reducing Cd content in the rice grain. The grain Cd content of the OsNramp5 knockout mutants was significantly lower than that of the wild-type rice T5105. However, the OsNramp5 knockout mutants still had much higher grain Cd content than the similar OsNramp5 mutants reported previously or the OsHAM3 overexpression line developed in our previous study. Pyramiding the OsNramp5 mutant allele and the OsHAM3 transgene in a double homozygous line could not further reduce grain Cd content. The OsNramp5 gene in T5105 has a haplotype II promoter, and its knockout mutation partially impairs Mn uptake in rice. Our results demonstrate that OsHMA3 overexpression works more efficiently in generating low-Cd rice grain than OsNramp5 knockout mutation without affecting Mn uptake in rice.

